# SVbyEye: A visual tool to characterize structural variation among whole-genome assemblies

**DOI:** 10.1101/2024.09.11.612418

**Authors:** David Porubsky, Xavi Guitart, DongAhn Yoo, Philip C. Dishuck, William T. Harvey, Evan E. Eichler

## Abstract

**Motivation:** We are now in the era of being able to routinely generate highly contiguous (near telomere-to-telomere) genome assemblies of human and nonhuman species. Complex structural variation and regions of rapid evolutionary turnover are being discovered for the first time. Thus, efficient and informative visualization tools are needed to evaluate and directly observe structural differences between two or more genomes.

**Results:** We developed SVbyEye, an open-source R package to visualize and annotate sequence-to-sequence alignments along with various functionalities to process alignments in PAF format. The tool facilitates the characterization of complex structural variants in the context of sequence homology helping resolve the mechanisms underlying their formation.

**Availability and implementation:** SVbyEye is available at https://github.com/daewoooo/SVbyEye.

## Introduction

Informative and efficient visualization of genomic structural variation (SV) is an important step to evaluate the validity of the most complex regions of the genome, helping us to develop new hypotheses and draw biological conclusions. With advances in long-read sequencing technologies, such as HiFi (high-fidelity) PacBio (Wenger et al. 2019) and ONT (Oxford Nanopore Technologies) (Deamer and Branton 2002), we are now able to fully assemble even the most complex regions of the genome, such as segmental duplications (Vollger et al. 2022), acrocentric regions (Guarracino et al. 2023), and centromeres (Logsdon et al. 2024) into continuous, highly accurate linear assemblies—also known as telomere-to-telomere (T2T) assemblies (Jarvis et al. 2022; Nurk et al. 2022). A large part of our understanding of the evolution of complex biological systems comes from comparative analyses, including direct visual observations (Paparella et al. 2023; Yoo et al. 2024).

The challenge with these analyses is that many large-scale structural changes between genomes are mediated by large, highly identical repeat sequences that are not readily annotated by existing software. This necessitates the development of visualization tools to complement T2T comparative studies. We developed SVbyEye for three purposes: 1) to directly characterize structurally complex regions, including insertions, duplications, deletions and inversions, by comparison to a linear genome reference; 2) to place these changes in the context of sequence homology by characterizing associated sequence identity; and 3) to define the breakpoints, including the length and orientation of homologous sequence mediating the rearrangement. SVbyEye is inspired by the previously developed tool called Miropeats (Parsons 1995) and brings its visuals to the popular scripting language R and visualization paradigm using ggplot2 (Wickham 2009).

## Materials and Methods

SVbyEye uses as input DNA sequence alignments in PAF (Pairwise mApping Format) format, which can be easily generated with minimap2 (Li 2018). In principle, however, any sequence-to-sequence aligner that can export alignments in standard PAF format should be sufficient. We note, however, that we tested our tool only using minimap2 alignments. Such alignments can be read using the ‘readPaf’ function. Subsequently, imported alignments can be filtered and flipped into the desired orientation using ‘filterPaf’ and ‘flipPaf’ functions, respectively,

### Visualization modes

There are four visualization modes offered by SVbyEye: visualization of pairwise alignments, alignments between more than two sequences, alignments within a single sequence, and whole-genome alignments.

The main function of this package, called ‘plotMiro’, serves to visualize pairwise sequence alignments in a horizontal layout with the target (reference) sequence at the top and the query at the bottom (**Fig. 1A**). The user has control over a number of visual and alignment processing features. For instance, users can color sequence alignments by their orientation or percentage of matched bases (**Supplementary Notes**).

**Figure 1:**
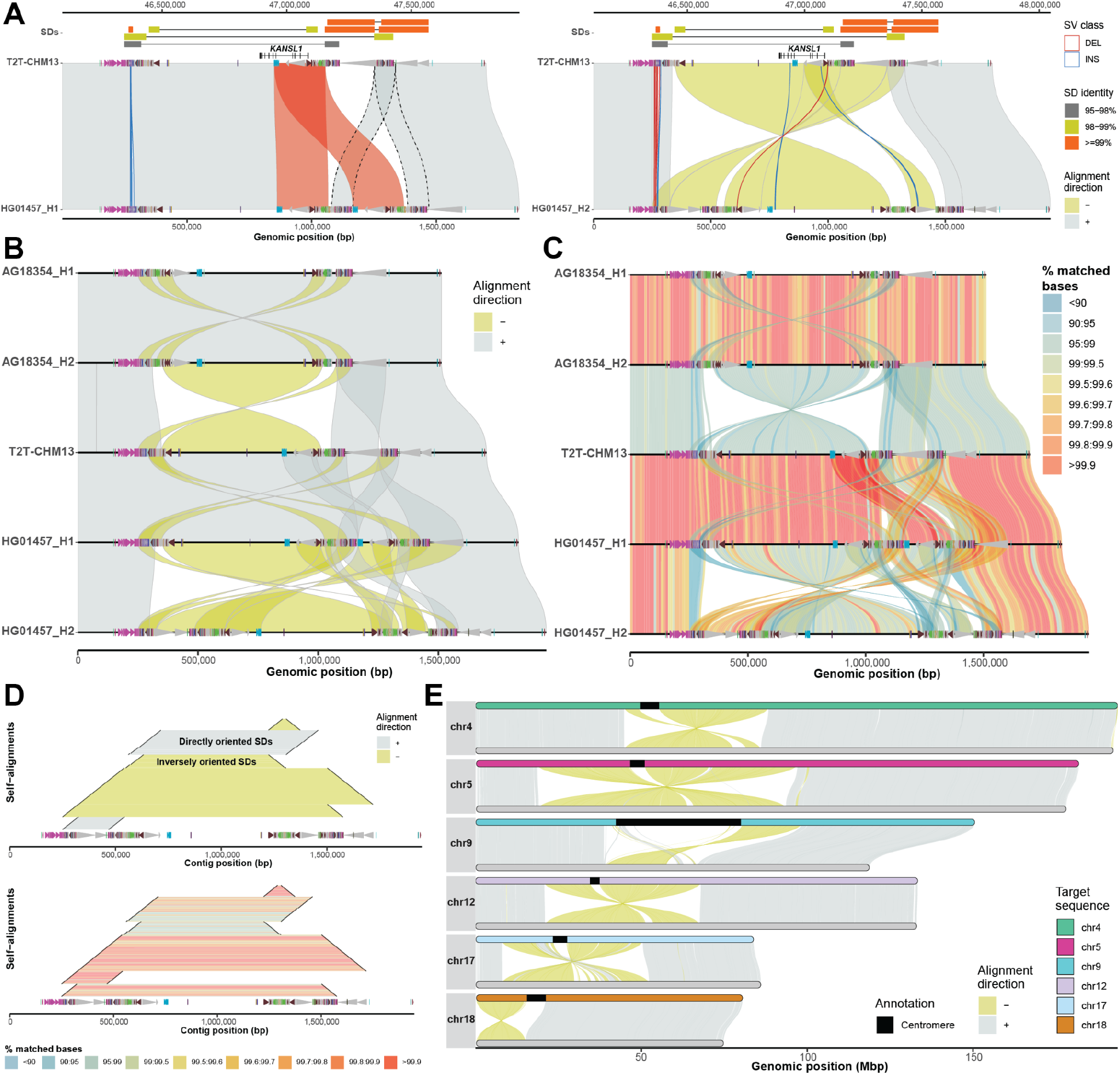
Example of SVbyEye visualization modes. **A**) The plot depicts a minimap alignment of a 1.7 Mbp region from chromosome 17q21.31 of two human sequences: HG01457 haplotypes (query) vs. T2T-CHM13 reference (target). Segmental duplication (SD) pairwise alignments are shown (top) (connected by horizontal line) colored by their sequence identity with gene annotation (*KANSL1* exons) depicted below as annotated in the UCSC Genome Browser. Minimap2 alignments are shown as gray (forward ‘+’ orientation) and yellow (reverse ‘-’ orientation) polygons between query (bottom) and target (top) sequences. Duplicon annotations as defined by DupMasker (Jiang et al. 2008) are indicated for both query and target sequences by colored arrowheads pointing forward or backward based on their orientation. An SV embedded within the SDs between query and target sequences (≥1 kbp) is highlighted as blue (insertion) and red (deletion) outlines facilitating breakpoint definition. **B**) A “stacked” SVbyEye plot depicting the 17q21.31 region for two chimpanzee haplotypes followed by three human haplotypes from T2T-CHM13 and HG01457. Each sequence is compared to the sequence immediately above and clearly defines a 750 kbp inversion between chimpanzee and human flanked by inverted repeats. A larger 900 kbp inversion polymorphism is also identified in human mediated by inverted SDs. **C**) The plot shows the same alignments as in B but with a “% identity grid” colored by the percentage of matched bases per 10-kbp-long sequence bin. Human inversion shows significant divergence indicating a deeper coalescence of the 17q21.31 region (Zody et al. 2008). **D**) A ‘horizontal dotplot’ visualization that shows self-alignments of HG01457 (haplotype 2) indicating the size (black line), the orientation (inverted=yellow and gray=direct; top panel), and the pairwise identity (colored grid; bottom panel). The largest and most identical segments are preferred sites for non-allelic homologous recombination (NAHR) breakpoints. **E**) A T2T view of six chimpanzee chromosomes (query, bottom) aligned to human syntenic chromosomes (T2T-CHM13, target, top). This view readily defines the extent of paracentric and pericentric inversions.

SVbyEye also allows visualization of alignments between more than two sequences. This can be done by aligning multiple sequences to each other using so-called all-versus-all (AVA) or stacked alignments and submitting them to the ‘plotAVA’ function. In this way, alignments are visualized in subsequent order where the alignment of the first sequence is shown with respect to the second and then second sequence to the third and so on (**Fig. 1B**). Many of the same parameters from ‘plotMiro’ also apply to ‘plotAVA’ as well. We illustrate a use of binning PAF alignments into defined bins (by setting a parameter ‘binsize’) and coloring them by the percentage of matched bases—a useful feature to reflect regional or pairwise differences in sequence identity (**Fig. 1C**).

To accommodate visualization of regions that are homologous to each other within a single sequence, we implemented the ‘plotSelf’ function. This function takes PAF alignments of a sequence to itself and visualizes them in a so-called horizontal dotplot (**Fig. 1D**). Such visualization can tell us a relative orientation, identity, and size of intrachromosomally aligned regions, an important feature of segmental duplications that predispose intervening sequence to recurrent rearrangements (Itsara et al. 2012; Coe et al. 2014; Porubsky et al. 2022). We note that self-alignments can also be visualized as arcs or arrowed rectangles connecting aligned regions (**Supplementary Notes**).

To allow for a full overview of whole-genome assembly with respect to a reference, SVbyEye offers ‘plotGenome’ function. With this function whole-genome alignments can be visualized to observe large structural rearrangements, such as large para- and pericentromeric inversions between the chimpanzee and human genomes (**Fig. 1E**).

### Alignment processing and annotation functionalities

SVbyEye has the ability to break PAF alignments at the positions of insertions and deletions and thereby delineate their breakpoints. This is done by parsing alignment CIGAR strings if reported in the PAF file. Thus, by setting the minimum size of insertions and deletions to be reported, one can visualize SVs as red (deletions) and blue (insertions) outlines (**Fig. 1A**). For further interrogation users can also opt to report alignments embedded insertions and/or deletions in a data table format using the ‘breakPaf’ function.

An important feature of SVbyEye is its capability to annotate query and target sequences with genomic ranges such as gene position, position of segmental duplications, or other DNA functional elements. This is done by adding extra annotation layers on top of the target and/or query alignments using the ‘addAnnotation’ function (**Supplementary Notes**). Annotation ranges are visualized as either a rectangle or an arrowhead. Arrowheads are especially useful for conveying an orientation of a genomic range. Similar to PAF alignments, annotation ranges can also be colored by a user defined color scheme (**Fig. 1A**). If there is a need to highlight specific PAF alignments between a query and a target, one can do so with the ‘add Alignments’ function that adds selected alignment(s) over the original plot highlighted by a unique outline and/or color (**Fig. 1A**).

There are several other useful functionalities that come with SVbyEye, for instance, lifting coordinates from target to query and vice versa provided by the ‘lift Ranges To Alignment’ function. Users can also subset alignments from a desired region on a target sequence using the ‘subsetPaf’ function. Lastly, there is a possibility to disjoin PAF alignments at regions where two and more alignments overlap each other with the ‘disjoin Paf Alignments’ function to provide exact boundaries of duplicated regions (**Fig. 1A**).

## Conclusion

We developed SVbyEye, a data visualization R package, to facilitate direct observation of structural differences between two or more sequences. SVbyEye provides several visualization modes depending on the desired application. It offers ample ways to annotate both query and target sequences along with many functionalities to process alignments in PAF format. A more detailed package documentation along with code examples can be found at: https://htmlpreview.github.io/?https://github.com/daewoooo/SVbyEye/blob/master/man/doc/SVbyEye.html.

## Acknowledgements

This article is subject to HHMI’s Open Access to Publications policy. HHMI lab heads have previously granted a non-exclusive CC BY 4.0 license to the public and a sublicensable license to HHMI in their research articles. Pursuant to those licenses, the author-accepted manuscript of this article can be made freely available under a CC BY 4.0 license immediately upon publication.

