## Supplementary Notes for "SVbyEye: A visual tool to characterize structural variation among whole-genome assemblies"

***David Porubsky***

**2024-09-17**

**Package**

SVbyEye 0.99.0

### 1 Introduction

Informative and efficient visualization of genomic structural variation (SV) is an important step to evaluate structurally complex regions of the genome and help us to begin drawing biological conclusions. With advances in long-read sequencing technologies, such as HiFi (high-fidelity) PacBio (Wenger et al. 2019) and ONT (Oxford Nanopore Technologies) (Deamer and Branton 2002), we are now able to fully assemble even the most complex regions of the genome. Thus, efficient and informative visualization tools are needed to evaluate and directly observe structural differences between two or more genomes. We have developed SVbyEye exactly for this purpose such that we are able to directly observe complexity of a human (or other organism) genome in question with respect to a linear genome reference. SVbyEye is inspired by the previously developed tool Miropeats (Parsons 1995) and brings its visuals to the popular scripting language R and visualization paradigm using ggplot2 (Wickham 2016).

### 2 Generation of input alignments in PAF format

In order to create input alignments for SVbyEye visualization, we usually use minimap2 (version 2.24 or higher) (Li 2016); however, any sequence-to-sequence aligner that can export alignments in PAF format should be sufficient. We note, however, that we tested our tool only using minimap2 alignments.

When running minimap2 alignments ensure that parameter -a is not set because it would export alignment in SAM format. Also make sure that parameter -c is set in order to export CIGAR strings as 'cg' tag in the output PAF. Lastly, consider setting the parameter '–eqx' in order to output CIGAR string with '=/X' operators to define sequence match/mismatch, respectively.

#### 2.1 Generate assembly-to-reference minimap alignments

minimap2 -x asm20 -c –eqx –secondary=no {reference.fasta} {query.fasta} > {output.alignment}

#### 2.2 Generate sequence-to-sequence minimap alignments

minimap2 -x asm20 -c –eqx –secondary=no {target.fasta} {query.fasta} > {output.alignment}

#### 2.3 Generate all-versus-all minimap alignments

minimap2 -x asm20 -c –eqx -D -P –dual=no {input.multi.fasta} {input.multi.fasta} > {output.ava.alignment}

### 3 Reading and filtering alignments in PAF format

The *SVbyEye* package expects sequence alignments in PAF format as input. Such alignments can be read using the `readPaf()` function. Subsequently, such alignments can be filtered using the `filterPaf()` function. Lastly, one can also orient PAF alignments based on the desired majority orientation (`majority.strand = '+' or '-'`) using the `flipPaf()` function. With this function, the user can flip all alignments in the PAF file to opposite orientation with parameter `force = TRUE`.

First load the *SVbyEye* package.

```
## Load the SVbyEye package
library(SVbyEye)
```

```
## Get PAF file to read
paf.file <- system.file("extdata", "test1.paf",
    package = "SVbyEye"
)
## Read in PAF
paf.table <- readPaf(
    paf.file = paf.file,
    include.paf.tags = TRUE, restrict.paf.tags = "cg"
)
```

```
## Example PAF table, SVbyEye expects given column names
paf.table
## # A tibble: 4 x 13
##   q.name        q.len q.start  q.end strand t.name  t.len t.start  t.end n.match
##   <chr>         <dbl>   <dbl>  <dbl> <chr>  <chr>   <dbl>   <dbl>  <dbl>   <dbl>
## 1 query.region 2.19e7  1.64e7 1.66e7 +      targe~ 8.33e7  1.92e7 1.93e7  161374
## 2 query.region 2.19e7  1.61e7 1.62e7 +      targe~ 8.33e7  1.89e7 1.90e7   90136
## 3 query.region 2.19e7  1.62e7 1.63e7 +      targe~ 8.33e7  1.90e7 1.91e7   49613
## 4 query.region 2.19e7  1.63e7 1.64e7 -      targe~ 8.33e7  1.91e7 1.92e7  106633
## # i 3 more variables: aln.len <dbl>, mapq <dbl>, cg <chr>

## Filter alignment based on size
filterPaf(paf.table = paf.table, min.align.len = 100000)
## # A tibble: 2 x 13
##   q.name        q.len q.start  q.end strand t.name  t.len t.start  t.end n.match
##   <chr>         <dbl>   <dbl>  <dbl> <chr>  <chr>   <dbl>   <dbl>  <dbl>   <dbl>
## 1 query.region 2.19e7  1.64e7 1.66e7 +      targe~ 8.33e7  1.92e7 1.93e7  161374
## 2 query.region 2.19e7  1.63e7 1.64e7 -      targe~ 8.33e7  1.91e7 1.92e7  106633
## # i 3 more variables: aln.len <dbl>, mapq <dbl>, cg <chr>

## Force to flip orientation of PAF alignments
flipPaf(paf.table = paf.table, force = TRUE)
## # A tibble: 4 x 16
##   q.name        q.len q.start  q.end strand t.name  t.len t.start  t.end n.match
## * <chr>         <dbl>   <dbl>  <dbl> <chr>  <chr>   <dbl>   <dbl>  <dbl>   <dbl>
## 1 query.region 2.19e7 5384372 5.55e6 -      targe~ 8.33e7  1.92e7 1.93e7  161374
## 2 query.region 2.19e7 5703733 5.79e6 -      targe~ 8.33e7  1.89e7 1.90e7   90136
## 3 query.region 2.19e7 5652689 5.70e6 -      targe~ 8.33e7  1.90e7 1.91e7   49613
## 4 query.region 2.19e7 5545965 5.65e6 +      targe~ 8.33e7  1.91e7 1.92e7  106633
## # i 6 more variables: aln.len <dbl>, mapq <dbl>, cg <chr>, query.flip <lgl>,
## #   seq.pair <chr>, target.flip <lgl>
```

### 4 Visualization of sequence-to-sequence alignments

The main function of this package is `plotMiro()` and it can be used for visualization of a single sequence aligned to the reference or more than two sequences aligned to each other. SVbyEye is visualizing such alignments in a horizontal layout with the target sequence being on at the top and the query at the bottom.

To create a simple Miropeats style plot, follow thevinstructions below.

```
## Get PAF to plot
paf.file <- system.file("extdata", "test1.paf",
    package = "SVbyEye"
)
## Read in PAF
paf.table <- readPaf(
    paf.file = paf.file,
    include.paf.tags = TRUE, restrict.paf.tags = "cg"
)
```

```r
## Make a plot colored by alignment directionality
plotMiro(paf.table = paf.table, color.by = "direction")
```

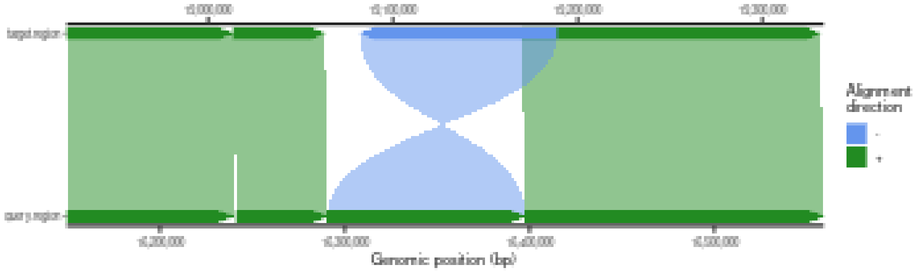

Users have control over a number of visual features that are documented within a `plotMiro()` function. Function documentation can be reported by `?plotMiro`.

For instance, users can specify their own color scheme to designate alignments in plus (forward) or minus (reverse) direction.

```r
## Use custom color palette to color alignment directionality
plotMiro(paf.table = paf.table,
         color.palette = c("+" = "azure3", "-" = "yellow3"))
```

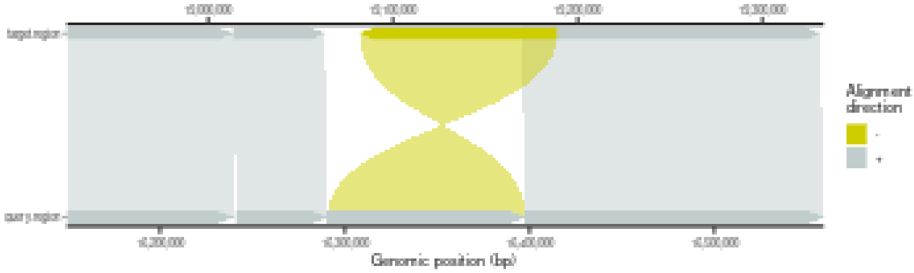

Next, users can decide to color alignments by their identity instead of direction.

```r
## Color alignments by sequence identity
plotMiro(paf.table = paf.table, color.by = "identity")
```

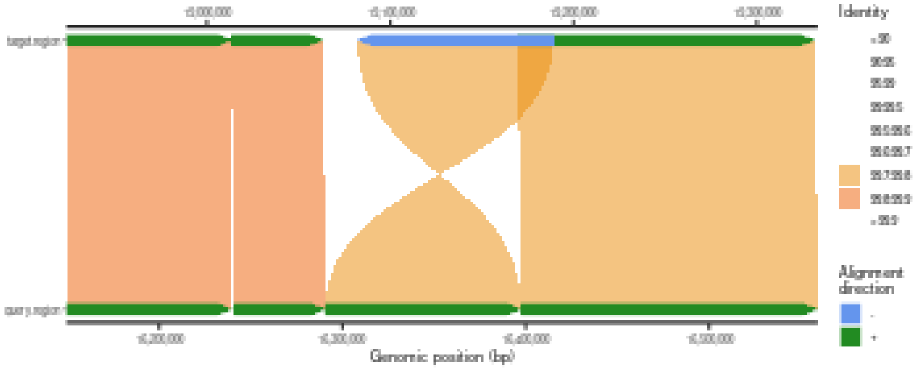

We advise that coloring alignments by sequence identity is the most helpful in connection with binning the alignments into separate bins. This is useful when investigating regions of high and low sequence identity between the target and query sequences. We note that when the parameter 'binsize' is defined, the parameter 'color.by' is set to 'identity' by default.

```r
## Plot binned alignments
plotMiro(paf.table = paf.table, binsize = 1000)
```

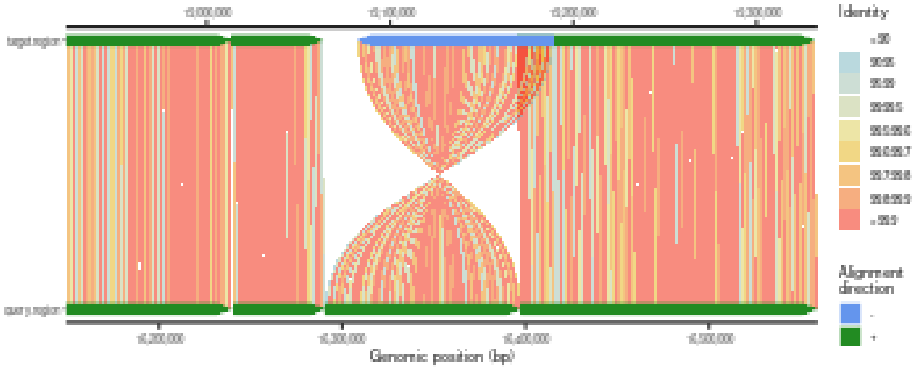

Currently there are preset breaks of sequence identity (see `?plotMiro`) that can be changed by setting the parameter 'perc.identity.breaks'.

#### 4.1 Adding annotations to the Miropeats style plot

Often certain regions need to marked between the query and target alignments such as gene position, position of segmental duplications (SDs), or other DNA functional elements. This can be done by adding extra layers of annotation ranges either on top of the target or query alignments. Annotation ranges are expected to be submitted to the `addAnnotation()` function as a GenomicRanges object and are visualized as either an arrowhead (can reflect range orientation) or rectangle.

```r
## Make a plot
plt <- plotMiro(paf.table = paf.table)
## Load target annotation file
target.annot <- system.file("extdata", "test1_target_annot.txt",
                            package = "SVbyEye")
target.annot.df <- read.table(target.annot, header = TRUE, sep = "\t",
                              stringsAsFactors = FALSE)
target.annot.gr <- GenomicRanges::makeGRangesFromDataFrame(target.annot.df)
## Add target annotation as arrowhead
addAnnotation(ggplot.obj = plt,
              annot.gr = target.annot.gr, coordinate.space = "target")
```

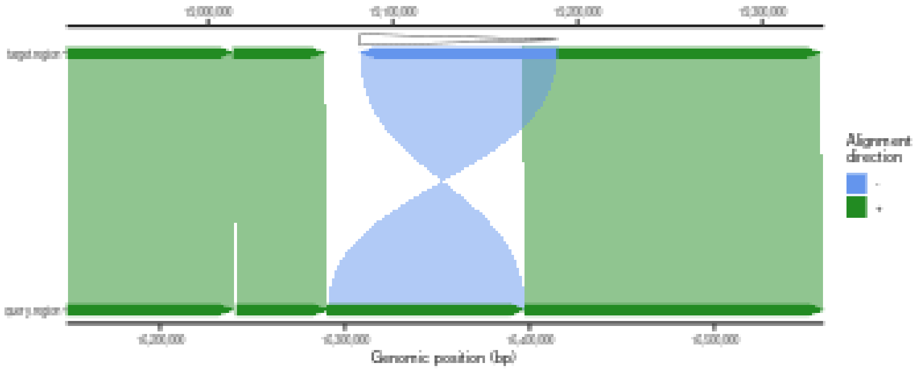

```r
## Load query annotation file
query.annot <- system.file("extdata", "test1_query_annot.txt",
                           package = "SVbyEye")
query.annot.df <- read.table(query.annot, header = TRUE, sep = "\t",
                             stringsAsFactors = FALSE)
query.annot.gr <- GenomicRanges::makeGRangesFromDataFrame(query.annot.df)
## Add query annotation as rectangle
addAnnotation(ggplot.obj = plt,
              annot.gr = query.annot.gr, shape = "rectangle",
              coordinate.space = "query")
```

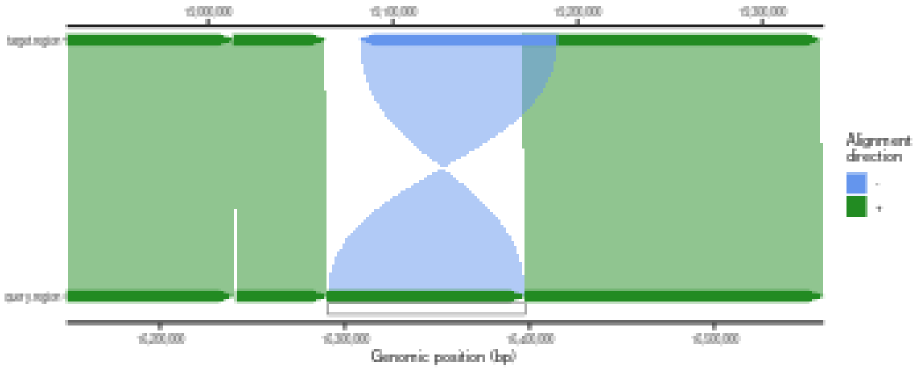

SVbyEye also offers extra functionalities to lift between query and target coordinates and vice versa based on the underlying PAF alignment file.

```r
## Lift target annotation to query and plot
target.gr <- as("target.region:19100000-19150000", "GRanges")
lifted.annot.gr <- liftRangesToAlignment(paf.table = paf.table,
                                         gr = target.gr,
                                         direction = "target2query")
plt1 <- addAnnotation(
    ggplot.obj = plt, annot.gr = target.gr, shape = "rectangle",
     coordinate.space = "target"
 )
 addAnnotation(
     ggplot.obj = plt1, annot.gr = lifted.annot.gr, shape = "rectangle",
     coordinate.space = "query"
 )
```

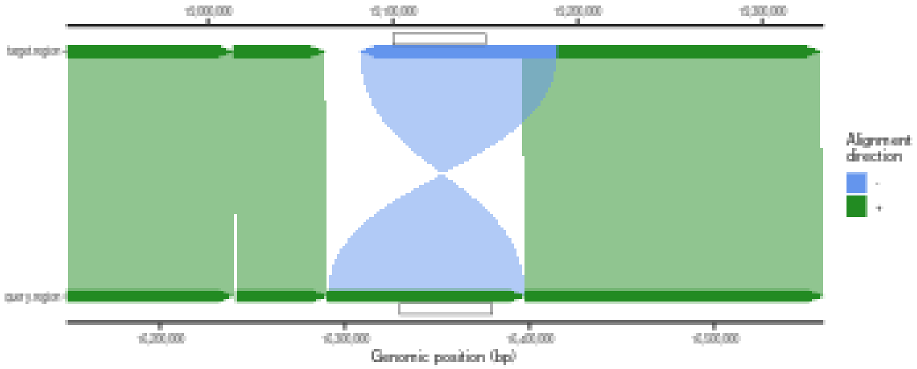

For instance, inversions are often flanked by long segments of highly identical SDs. These can be visualized as follows. The direction of these segments can be reflected as well in case the strand is defined in the GenomicRanges object.

```
## Load segmental duplication annotation
sd.annot <- system.file("extdata", "test1.sd.annot.RData", package = "SVbyEye")
sd.annot.gr <- get(load(sd.annot))
## Create a custom discrete levels based on
sd.categ <- findInterval(sd.annot.gr$fracMatch, vec = c(0.95, 0.98, 0.99))
sd.categ <- dplyr::recode(sd.categ,
                          "0" = "<95%", "1" = "95-98%",
                          "2" = "98-99%", "3" = ">=99%")
sd.categ <- factor(sd.categ, levels = c("<95%", "95-98%", "98-99%", ">=99%"))
sd.annot.gr$sd.categ <- sd.categ
## Define a custom color palette
color.palette <- c(
   "<95%" = "gray72", "95-98%" = "gray47", "98-99%" = "#cccc00",
   ">=99%" = "#ff6700"
)
## Add annotation to the plot
addAnnotation(
   ggplot.obj = plt, annot.gr = sd.annot.gr, fill.by = "sd.categ",
   color.palette = color.palette, coordinate.space = "target"
)
```

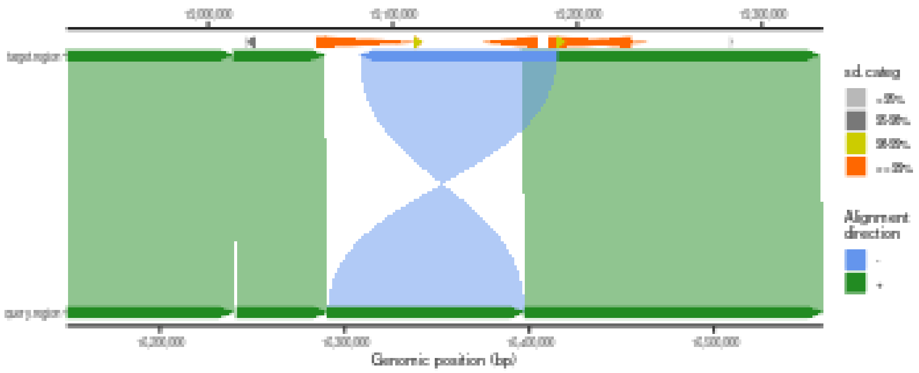

Importantly, each annotation layer can be given its own name to distinguish different layers of information added to the plot.

```
## Give an annotation layer its own label
## Add target annotation as arrowhead
plt1 <- addAnnotation(ggplot.obj = plt,
              annot.gr = target.annot.gr, coordinate.space = "target",
              annotation.label = 'Region')
## Add SD annotation
addAnnotation(
   ggplot.obj = plt1, annot.gr = sd.annot.gr, fill.by = "sd.categ",
   color.palette = color.palette, coordinate.space = "target",
   annotation.label = 'SDs'
)
```

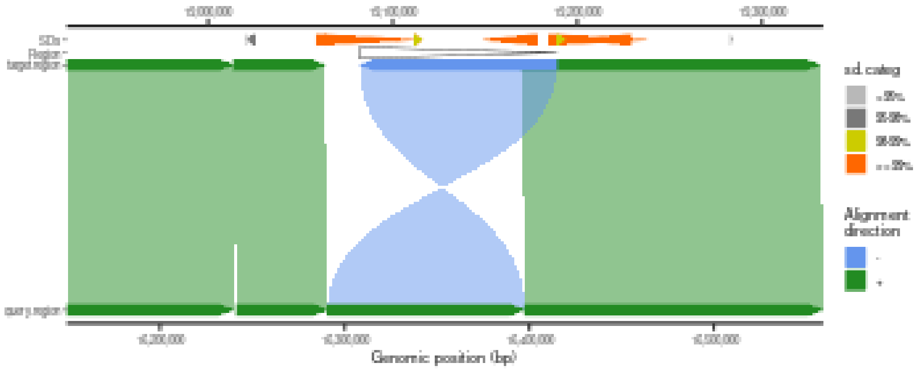

The user also has control over the level where an annotation track should appear. By default each annotation layer is displayed at the level defined by adding 0.05 fraction of the total size of y-axis in the target or query direction. This can of course be increased or decreased to zero, which means the annotation will be added at the same level as the target or query alignments.

```
## Add target annotation at the increased y-axis level
addAnnotation(ggplot.obj = plt, annot.gr = target.annot.gr,
              coordinate.space = "target", annotation.level = 0.2)
```

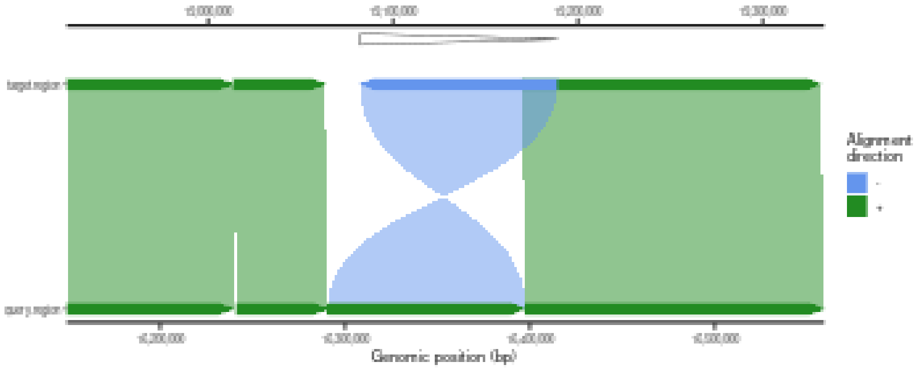

```
## Add target annotation at the zero level
addAnnotation(ggplot.obj = plt, annot.gr = target.annot.gr,
              coordinate.space = "target", annotation.level = 0)
```

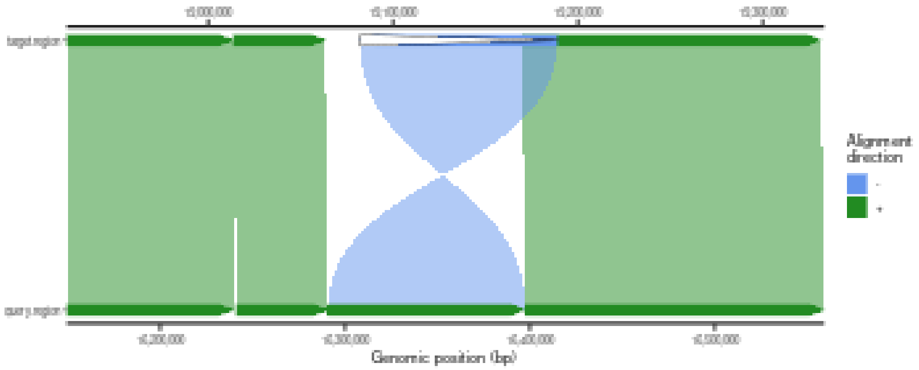

#### 4.2 Adding gene annotations

Gene annotation, such as individual exons or split genomic alignments, can be visualized as a set of genomic ranges that are connected by a horizontal line. This is defined based on a shared identifier (ID) that marks genomic ranges that are supposed to be grouped by such horizontal lines.

```
## Create gene-like annotation
test.gr <- GenomicRanges::GRanges(
              seqnames = 'target.region',
              ranges = IRanges::IRanges(start = c(19000000,19030000,19070000),
                                        end = c(19010000,19050000,19090000)),
                                        ID = 'gene1')
## Add single gene annotation
addAnnotation(ggplot.obj = plt, annot.gr = test.gr, coordinate.space = "target",
             shape = 'rectangle', annotation.group = 'ID', fill.by = 'ID')
```

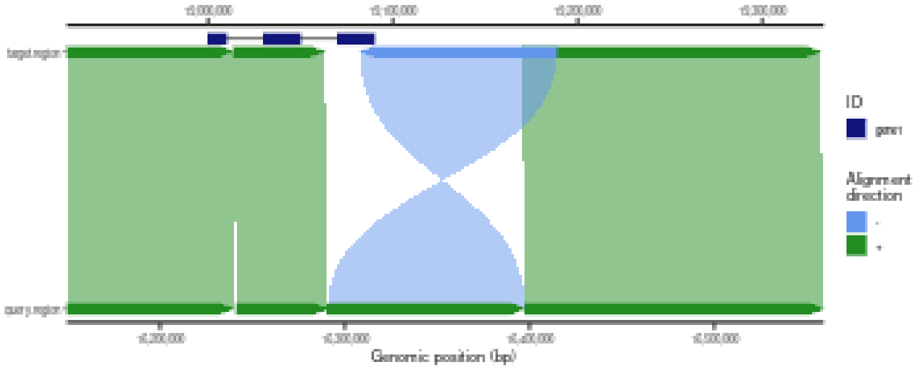

#### 4.3 Highlighting whole alignments

Sometimes there is a need to highlight the whole genomic alignment with a unique color. This can be done by overlaying the original alignment with an extra alignment.

```
## Make a plot
plt1 <- plotMiro(paf.table = paf.table, add.alignment.arrows = FALSE)
## Highlight alignment as a filled polygon
addAlignments(ggplot.obj = plt1, paf.table = paf.table[3,], fill.by = 'strand',
              fill.palette = c('+' = 'red'))
```

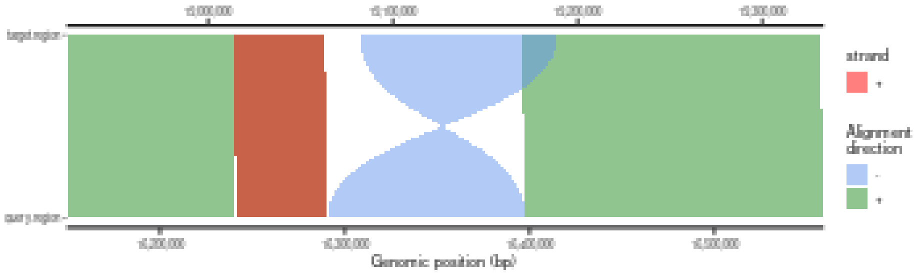

```
## Highlight alignment as outlined polygon by dashed line
addAlignments(ggplot.obj = plt1, paf.table = paf.table[4,], linetype = 'dashed')
```

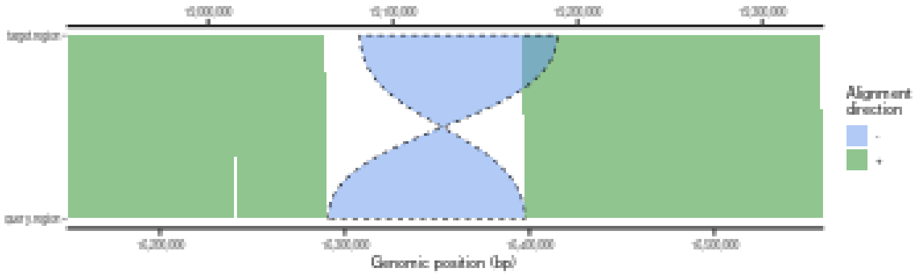

#### 4.4 Visualization of disjoint alignments

On many occasions alignments over the repetitive regions are redundant at positions of SDs. This results in overlapping alignments that often flank inverted regions. There is a function that allows breaking PAF alignments at these overlaps and reporting them as individual alignments.

```
## Disjoin PAF at target coordinates
disj.paf.table <- disjoinPafAlignments(paf.table = paf.table,
                                       coordinates = 'target')
## Plot disjoined alignments
plotMiro(paf.table = disj.paf.table, outline.alignments = TRUE)
```

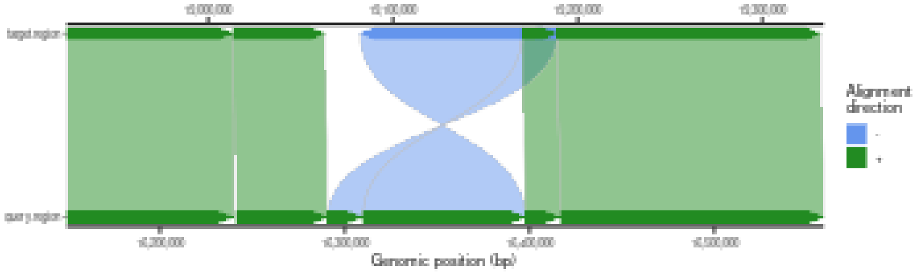

Disjoined alignments can also be highlighted as shown earlier.

```
## Plot disjoined alignments
plt <- plotMiro(paf.table = disj.paf.table, outline.alignments = TRUE,
                add.alignment.arrows = FALSE)
## Highlight disjoined alignments
addAlignments(ggplot.obj = plt, paf.table = disj.paf.table[c(4:5),],
              linetype = 'dashed')
```

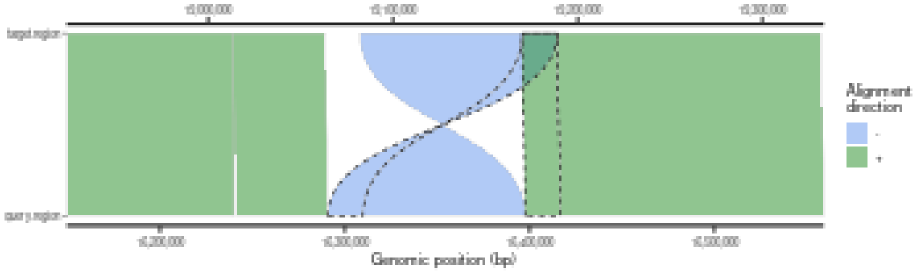

For convenience, PAF alignments can also be disjoined at user-defined region(s).

```
## Disjoin PAF at user defined coordinates
disj.gr <- GenomicRanges::GRanges(seqnames = 'query.region',
                                  ranges = IRanges::IRanges(start = 16300000,
                                                            end = 16350000))
disj.paf.table <- disjoinPafAlignments(paf.table = paf.table,
                                       coordinates = 'query',
                                       disjoin.gr = disj.gr)
## Plot disjoined alignments
plt <- plotMiro(paf.table = disj.paf.table, outline.alignments = TRUE)
## Highlight disjoined alignments
addAlignments(ggplot.obj = plt, paf.table = disj.paf.table[4,],
              linetype = 'dashed')
```

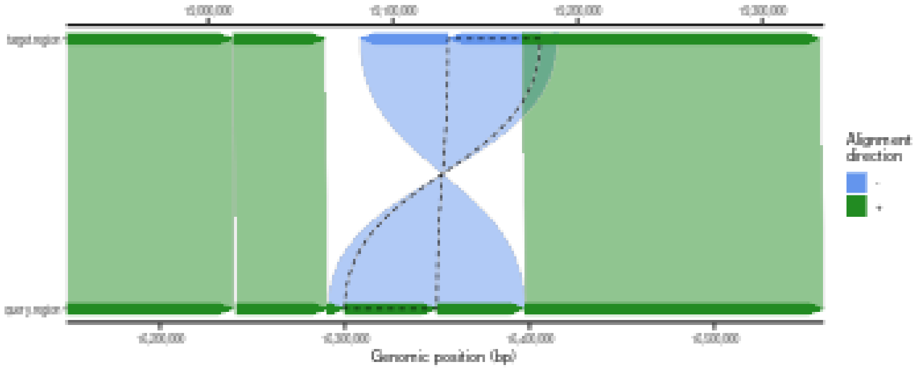

#### 4.5 Detection and visualization of insertions and deletions

Another important feature of SVbyEye is its ability to break PAF alignments at the positions of insertions and deletions. This can be done by setting the minimum size of the insertion and deletion to be reported and setting the way they are marked within the plot, either outlined or filled. By default, deletions are colored red and insertions are blue. Deletions and insertions are defined as sequences that are either missing or are inserted within a query sequence with respect to the target sequence.

```
## Load the data to plot
paf.file <- system.file("extdata", "test3.paf", package = "SVbyEye")
paf.table <- readPaf(paf.file = paf.file, include.paf.tags = TRUE,
                     restrict.paf.tags = "cg")

## Make plot and break alignment at the position where there are deletions >=50bp
plotMiro(paf.table = paf.table, min.deletion.size = 50, highlight.sv = "outline")
```

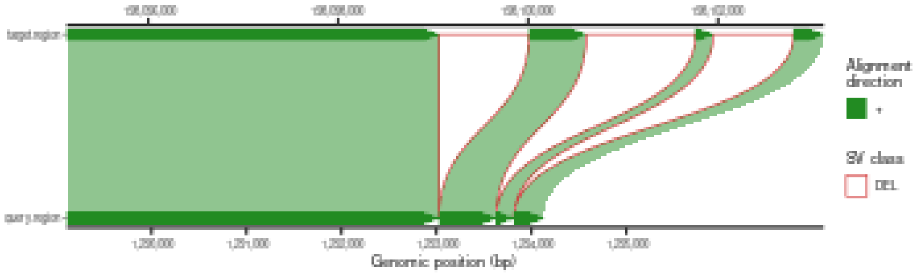

```
## Highlight detected deletion by filled polygons
plotMiro(paf.table = paf.table, min.deletion.size = 50, highlight.sv = "fill")
```

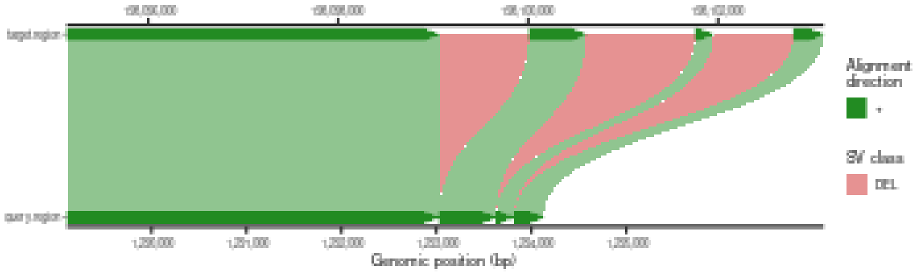

One can also opt to break the PAF alignments at insertions and/or deletions and just report them as a data table.

```
## Break PAF alignment at deletions >=50bp
alns <- breakPaf(paf.table = paf.table, min.deletion.size = 50)
## Print out detected deletions
alns$SVs
## # A tibble: 3 x 14
##   q.name        q.len q.start   q.end strand t.name   t.len t.start   t.end n.match
##   <chr>         <dbl>   <dbl>   <dbl> <chr>  <chr>    <dbl>   <dbl>   <dbl>   <dbl>
## 1 query.region 1.56e6 1293029 1.29e6 +      targe~ 1.98e8  1.98e8 1.98e8       0
## 2 query.region 1.56e6 1293632 1.29e6 +      targe~ 1.98e8  1.98e8 1.98e8       0
## 3 query.region 1.56e6 1293819 1.29e6 +      targe~ 1.98e8  1.98e8 1.98e8       0
## # i 4 more variables: aln.len <dbl>, mapq <dbl>, cg <chr>, aln.id <int>
```

#### 4.6 Narrowing alignment to a user-defined target region

There is a convenience function to narrow down a PAF alignment into a user-defined region by setting the desired target coordinates defined as a GenomicRanges object or a set of coordinates in this 'chromosome:start-end' format. With this, one can subset and cut PAF alignments at the exact target coordinates.

```
paf.file <- system.file("extdata", "test1.paf", package = "SVbyEye")
## Read in PAF
paf.table <- readPaf(paf.file = paf.file, include.paf.tags = TRUE,
                     restrict.paf.tags = "cg")
## Subset PAF
subsetPafAlignments(paf.table = paf.table,
                    target.region = "target.region:19050000-19200000")
## # A tibble: 3 x 13
##   q.name        q.len q.start   q.end strand t.name   t.len t.start   t.end n.match
##   <chr>         <dbl>   <dbl>   <dbl> <chr>  <chr>    <dbl>   <dbl>   <dbl>   <dbl>
## 1 query.region 2.19e7  1.63e7 1.64e7 -      targe~ 8.33e7  1.91e7 1.92e7  106633
## 2 query.region 2.19e7  1.64e7 1.64e7 +      targe~ 8.33e7  1.92e7 1.92e7   29958
## 3 query.region 2.19e7  1.63e7 1.63e7 +      targe~ 8.33e7  1.90e7 1.91e7   13183
## # i 3 more variables: aln.len <dbl>, mapq <dbl>, cg <chr>
```

This functionality can be directly used within the `plotMiro` function by setting the parameter 'target.region'.

#### 4.7 Summarizing sequence composition as annotation heatmaps

Lastly, it is possibile to evaluate the sequence composition of either the query or target sequence. The `fasta2nucleotideContent()` function can calculate the frequency of a specific sequence pattern or an overall frequency of nucleotides within user-defined sequence bins. Subsequently, such nucleotide frequencies can be added to the plot as a heatmap.

```
## Get PAF to plot
paf.file <- system.file("extdata", "test_getFASTA.paf",
    package = "SVbyEye"
)
## Read in PAF
paf.table <- readPaf(
    paf.file = paf.file,
    include.paf.tags = TRUE, restrict.paf.tags = "cg"
)
## Make a plot colored by alignment directionality
plt <- plotMiro(paf.table = paf.table, color.by = "direction")
## Get query sequence composition
fa.file <- paf.file <- system.file("extdata", "test_getFASTA_query.fasta",
    package = "SVbyEye"
)
## Calculate GC content per 5kbp bin
gc.content <- fasta2nucleotideContent(fasta.file = fa.file,
                                      binsize = 5000, nucleotide.content = 'GC')
## Add GC content as annotation heatmap
addAnnotation(ggplot.obj = plt, annot.gr = gc.content, shape = 'rectangle',
              fill.by = 'GC_nuc.freq', coordinate.space = 'query',
              annotation.label = 'GCcont')
```

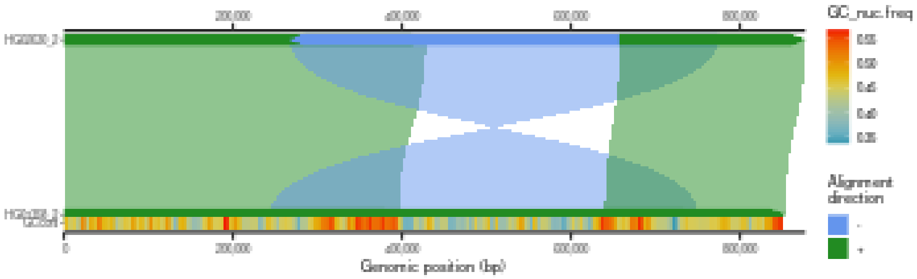

### 5 Visualization of all-versus-all sequence alignments

The SVbyEye also allows visualization of alignments between more than two sequences. This can be done by aligning multiple sequences to each other using so-called all-versus-all (AVA) or stacked alignments. In this way, alignments are visualized in subsequent order with alignments of the first sequence being shown with respect to the second and then second sequence to the third, etc. Alternatively, if the order of sequence in such a progressive alignment is known, one can avoid AVA alignment and align each sequence to the next in a defined order and merge these into a single PAF file. This is useful to visualize alignments between multiple sequences of the same or different species.

```r
## Get PAF to plot
paf.file <- system.file("extdata", "test_ava.paf", package = "SVbyEye")
## Read in PAF
paf.table <- readPaf(paf.file = paf.file, include.paf.tags = TRUE,
                     restrict.paf.tags = "cg")
## Make a plot colored by alignment directionality
plotAVA(paf.table = paf.table, color.by = "direction")
```

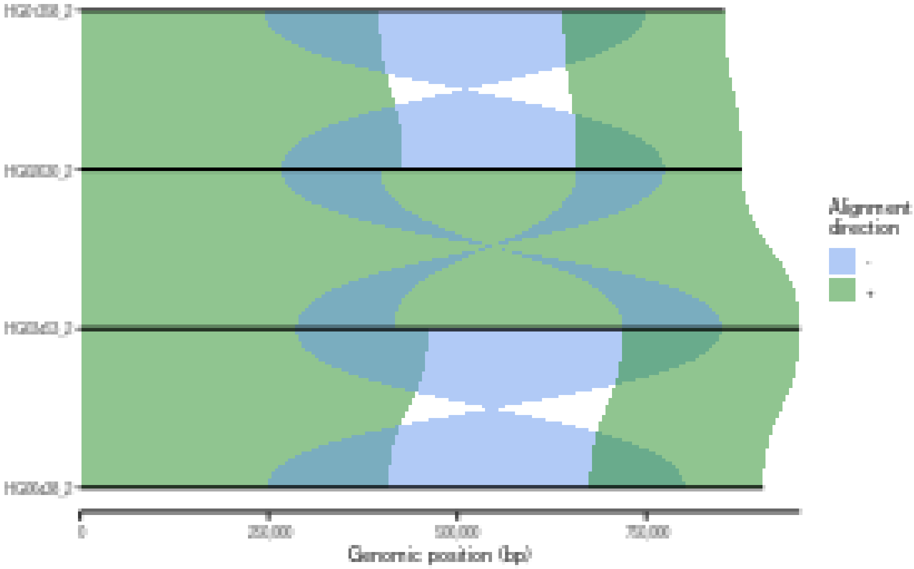

Many of the same parameter settings used for plotMiro apply to plotAVA as well. Briefly, users can color alignments based on their orientation or identity, define a desired color palette, and bin the alignments into user-defined bins.

```r
## Color by fraction of matched bases in each alignment
plotAVA(paf.table = paf.table, color.by = "identity",
        perc.identity.breaks = c(85, 90, 95))
```

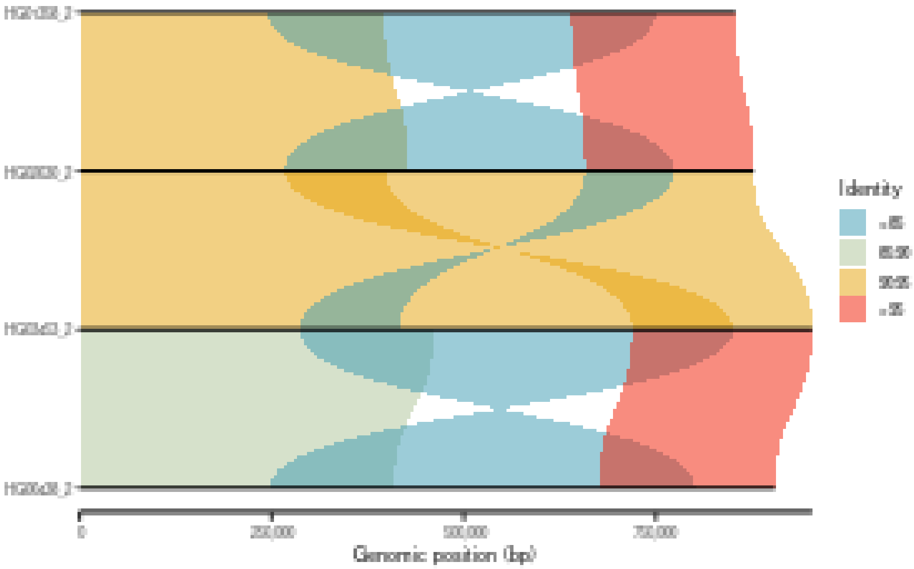

```r
## Use custom color palette to color alignment directionality
plotAVA(paf.table = paf.table, color.by = "direction",
        color.palette = c("+" = "azure3", "-" = "yellow3"))
```

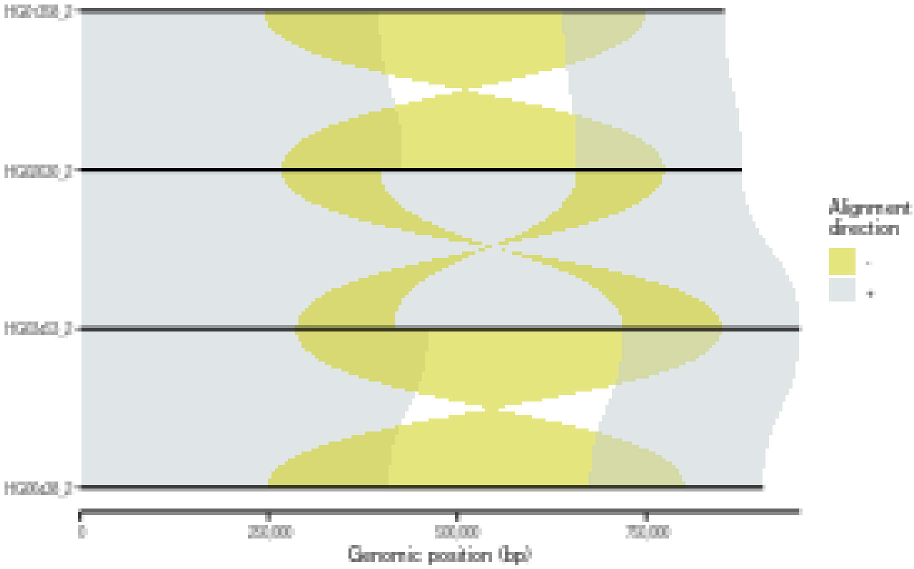

```r
## Bin PAF alignments into user-defined bin and color them by sequence identity
plotAVA(paf.table = paf.table, binsize = 20000)
```

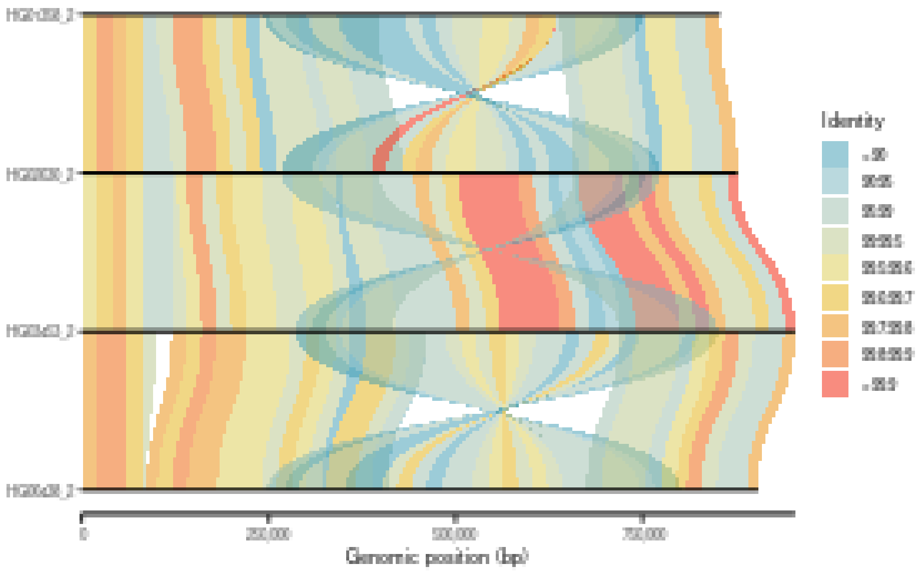

In addition to the above listed parameters, a user can also define the desired sequence order in which progressive alignments will be reported. Importantly, only samples defined in parameter ‘seqnames.order’ will be plotted.

```
## Define custom sample/sequence order
seqnames.order <- c("HG00438_2", "HG01358_2", "HG02630_2", "HG03453_2")
plotAVA(paf.table = paf.table, color.by = "direction",
        seqnames.order = seqnames.order)
```

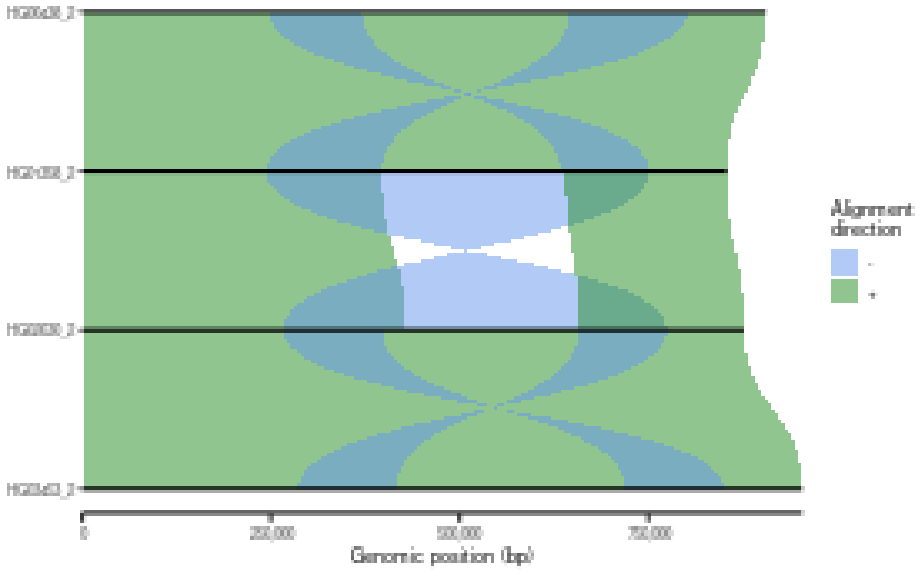

```
## Only samples present in custom sample order are being plotted
seqnames.order <- c("HG00438_2", "HG01358_2", "HG03453_2")
plotAVA(paf.table = paf.table, color.by = "direction",
        seqnames.order = seqnames.order)
```

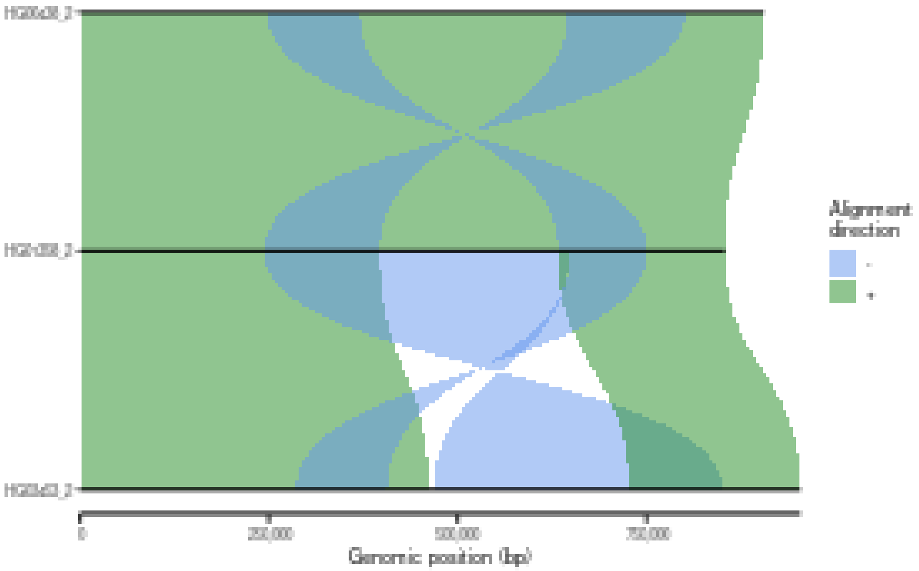

Adding annotation to AVA alignments differs slightly from what we learned with the plotMiro function. Since there are more than two sequences, annotation levels are defined based on the sequence identifier they belong to. This means users have to define an extra column that contains sequence identifiers to which each genomic range belongs to.

```
## Add annotation to all-versus-all alignments ##
plt <- plotAVA(paf.table = paf.table, color.by = 'direction')
annot.file <- system.file("extdata", "test_annot_ava.RData", package="SVbyEye")
annot.gr <- get(load(annot.file))
addAnnotation(ggplot.obj = plt, annot.gr = annot.gr, coordinate.space = 'self',
              y.label.id = 'ID')
```

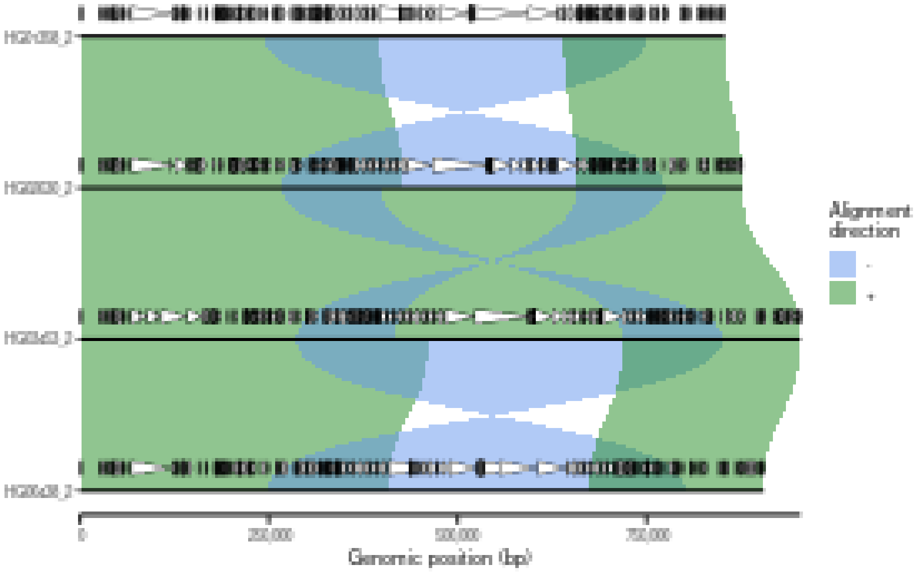

Again, users can set the annotation level to zero in order to plot each annotation directly on the line corresponding to each unique sequence/sample.

```
## Add annotation to the same level as the sequence/sample alignments
plt <- plotAVA(paf.table = paf.table, color.by = 'direction')
annot.file <- system.file("extdata", "test_annot_ava.RData", package="SVbyEye")
annot.gr <- get(load(annot.file))
addAnnotation(ggplot.obj = plt, annot.gr = annot.gr, coordinate.space = 'self',
              y.label.id = 'ID', annotation.level = 0)
```

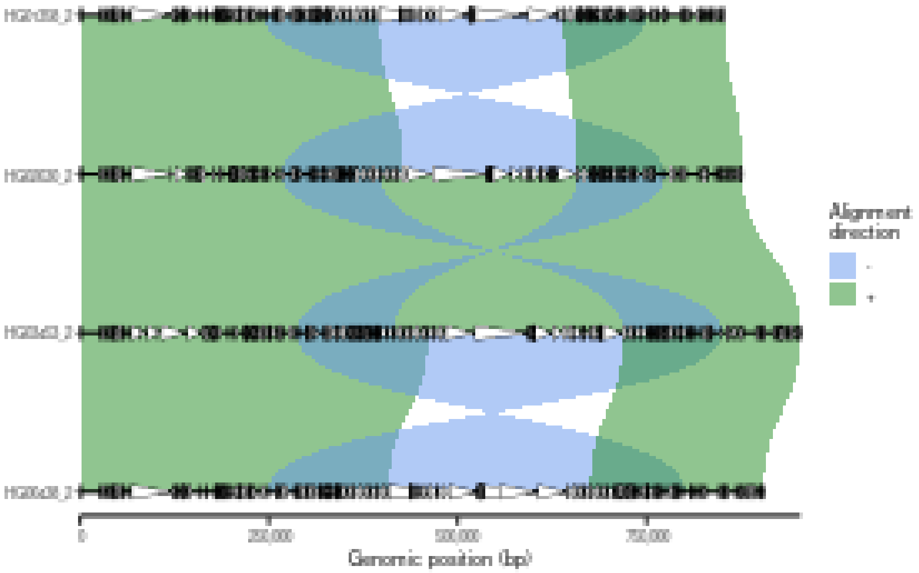

When plotting the annotation users can define a custom color palette that needs to be matched to the unique meta column in annotation ranges.

```
## Add annotation to the same level as the sequence/sample alignments
plt <- plotAVA(paf.table = paf.table, color.by = 'direction')
annot.file <- system.file("extdata", "test_annot_ava.RData", package="SVbyEye")
annot.gr <- get(load(annot.file))
## Define color palette
colors <- setNames(annot.gr$color, annot.gr$Repeat)
## Set fill.by variable to color each range using defined color palette
addAnnotation(ggplot.obj = plt, annot.gr = annot.gr, fill.by = 'Repeat',
              color.palette = colors, coordinate.space = 'self', y.label.id = 'ID',
              annotation.level = 0)
```

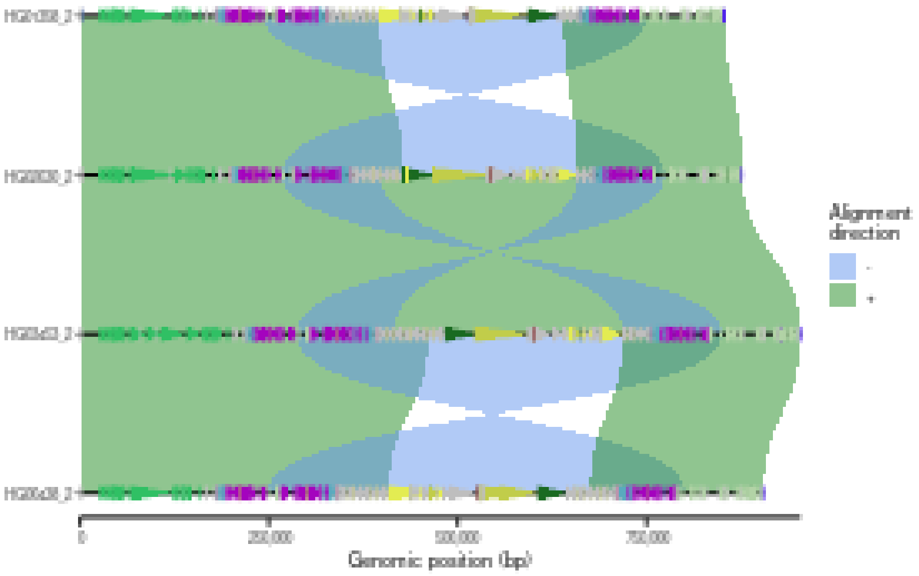

Lastly, there is again a possibility to break PAF alignments at the insertions and deletions and highlight them as outlined or filled polygons.

```
## Break PAF alignments at insertions and deletions and highlight them by an outline
plotAVA(paf.table = paf.table, color.by = "direction", min.deletion.size = 1000,
        min.insertion.size = 1000, color.palette = c("+" = "azure3", "-" = "yellow3"),
        highlight.sv = 'outline')
```

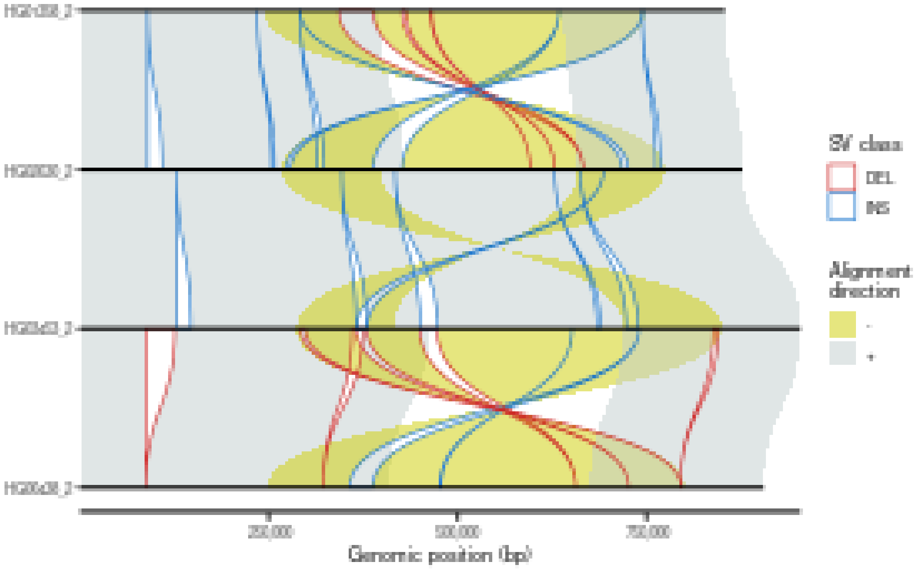

```
## Break PAF alignments at insertions and deletions and highlight them by filled polygons
plotAVA(paf.table = paf.table, color.by = "direction", min.deletion.size = 1000,
        min.insertion.size = 1000, color.palette = c("+" = "azure3", "-" = "yellow3"),
        highlight.sv = 'fill')
```

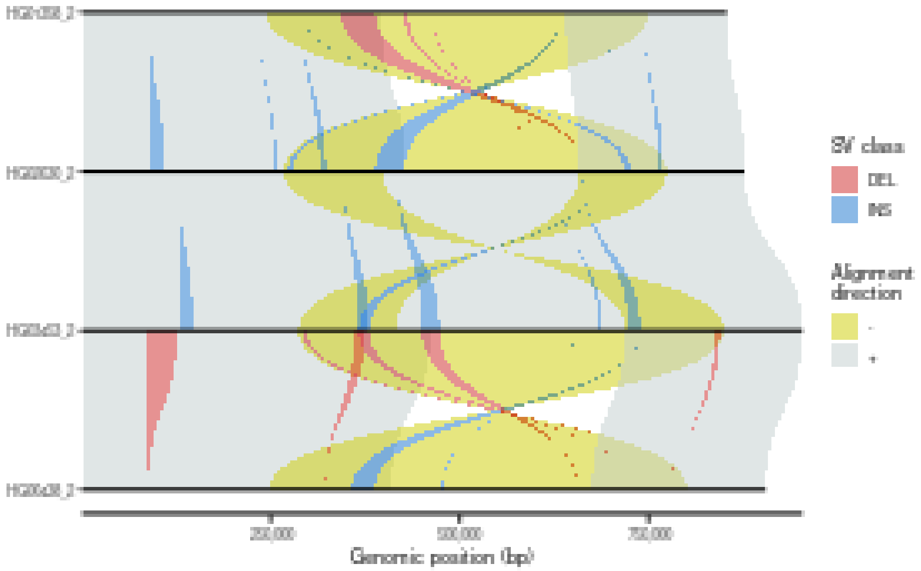

### 6 Visualization of self-alignments

There are cases when one wants to visualize regions that are homologous to each other within a single DNA sequence. Such regions can be reported by the alignment of a given sequence to itself. Such self-alignments are typical for SDs that are paralogous sequences with high sequence identity.

```
## Get PAF to plot
paf.file <- system.file("extdata", "test2.paf", package = "SVbyEye")
## Read in PAF
paf.table <- readPaf(paf.file = paf.file, include.paf.tags = TRUE,
                     restrict.paf.tags = "cg")
## Make a plot
## Plot alignment as horizontal dotplots and color by alignment directionality
plotSelf(paf.table = paf.table, color.by = "direction", shape = "segment")
```

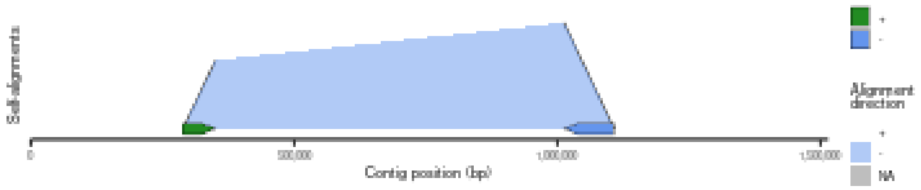

```
## Plot alignment as arcs and color by alignment directionality
plotSelf(paf.table = paf.table, color.by = "direction", shape = "arc")
```

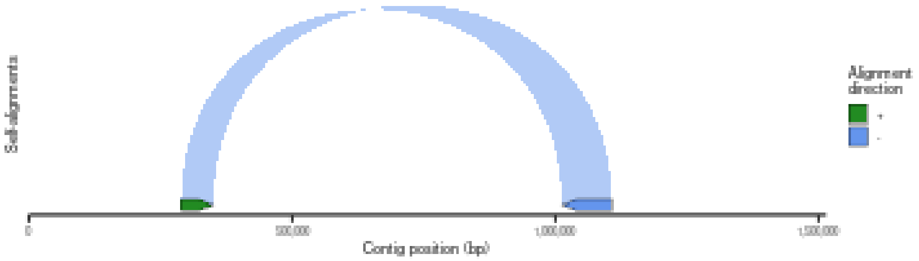

```r
## Plot alignment as arrows and color by alignment directionality
plotSelf(paf.table = paf.table, color.by = "direction", shape = "arrow")
```

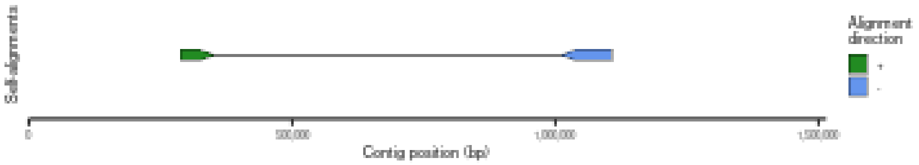

Again, some of the previous parameters apply to the self-alignments as well. For example, users can define the color palette and bin the alignments as well as break alignments at the insertions and deletions.

```r
## Plot alignment as arcs and color by alignment directionality
plotSelf(paf.table = paf.table, color.by = "direction", shape = "arc",
         color.palette = c("+" = "azure3", "-" = "yellow3"))
```

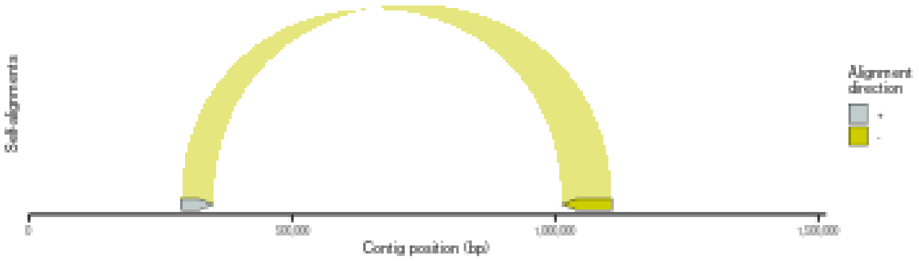

```r
## Bin PAF alignments into user-defined bin and color them by sequence identity
plotSelf(paf.table = paf.table, binsize = 1000)
```

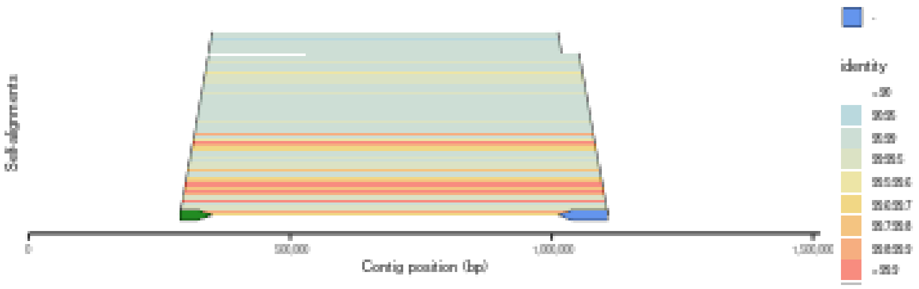

When breaking the self-alignments at insertions and deletions, proximal duplication is considered as a query and the distal duplication as a target. It means that sequence missing in the query is considered a deletion and sequence inserted in the query with respect to target sequence is considered an insertion.

```r
## Highlight structural variants within self-alignments
plotSelf(paf.table = paf.table, min.deletion.size = 50, min.insertion.size = 50,
         highlight.sv = "outline", shape = "arc")
```

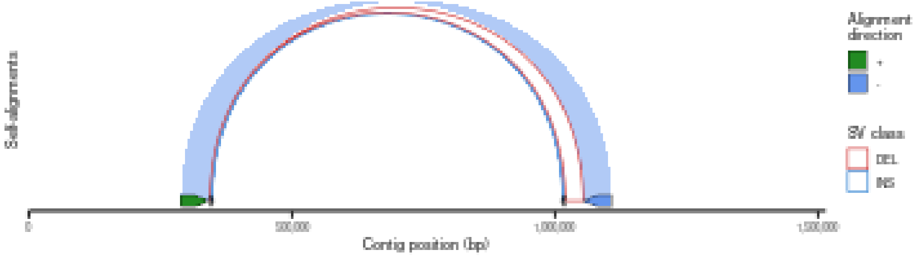

### 7 Visualization of whole-genome alignments

Lastly, SVbyEye allows plottin an overview of a whole-genome assembly or selected chromosomes with respect to a reference using the 'plotGenome' function. With this function whole-genome alignments can be visualized to observe large structural rearrangements with respect to a single reference.

```r
## Get PAF to plot ##
paf.file <- system.file("extdata", "PTR_test.paf", package = "SVbyEye")
## Read in PAF
paf.table <- readPaf(paf.file = paf.file)
## Make a plot ##
## Color by alignment directionality
plotGenome(paf.table = paf.table, chromosomes = paste0('chr', c(4, 5, 9, 12)),
           chromosome.bar.width = grid::unit(2, 'mm'),
           min.query.aligned.bp = 5000000)
```

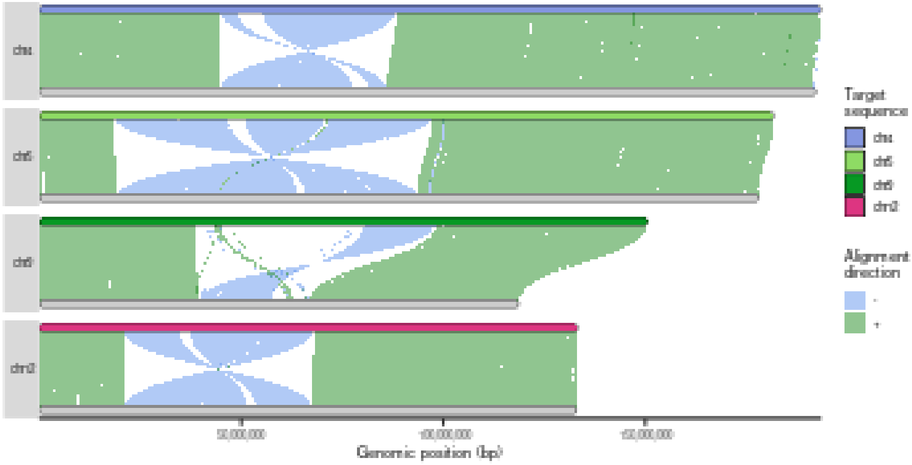

### A sessionInfo()

```
## R version 4.3.1 (2023-06-16)
## Platform: x86_64-pc-linux-gnu (64-bit)
## Running under: Ubuntu 22.04.2 LTS
##
## Matrix products: default
## BLAS:   /usr/lib/x86_64-linux-gnu/blas/libblas.so.3.10.0
## LAPACK: /usr/lib/x86_64-linux-gnu/lapack/liblapack.so.3.10.0
##
## locale:
##  [1] LC_CTYPE=en_US.UTF-8       LC_NUMERIC=C
##  [3] LC_TIME=en_US.UTF-8        LC_COLLATE=en_US.UTF-8
##  [5] LC_MONETARY=en_US.UTF-8    LC_MESSAGES=en_US.UTF-8
##  [7] LC_PAPER=en_US.UTF-8       LC_NAME=C
##  [9] LC_ADDRESS=C               LC_TELEPHONE=C
## [11] LC_MEASUREMENT=en_US.UTF-8 LC_IDENTIFICATION=C
##
## time zone: Europe/Bratislava
## tzcode source: system (glibc)
##
## attached base packages:
## [1] stats     graphics  grDevices utils     datasets  methods   base
##
## other attached packages:
## [1] SVbyEye_0.99.0   BiocStyle_2.28.0
##
## loaded via a namespace (and not attached):
##  [1] tidyselect_1.2.1            dplyr_1.1.4
##  [3] farver_2.1.2                Biostrings_2.70.3
##  [5] bitops_1.0-7                fastmap_1.1.1
##  [7] RCurl_1.98-1.14             tweenr_2.0.3
##  [9] GenomicAlignments_1.38.2    XML_3.99-0.16.1
## [11] digest_0.6.34               lifecycle_1.0.4
```

```
## [13] cluster_2.1.4                magrittr_2.0.3
## [15] compiler_4.3.1               rlang_1.1.3
## [17] tools_4.3.1                  utf8_1.2.4
## [19] yaml_2.3.8                   data.table_1.15.4
## [21] rtracklayer_1.62.0           knitr_1.43
## [23] labeling_0.4.3               S4Arrays_1.2.1
## [25] curl_5.2.1                   DelayedArray_0.28.0
## [27] abind_1.4-5                  BiocParallel_1.36.0
## [29] Rtsne_0.17                   withr_3.0.0
## [31] BiocGenerics_0.48.1          grid_4.3.1
## [33] polyclip_1.10-6              stats4_4.3.1
## [35] fansi_1.0.6                  colorspace_2.1-0
## [37] ggplot2_3.5.1                gtools_3.9.4
## [39] scales_1.3.0                 MASS_7.3-59
## [41] SummarizedExperiment_1.32.0 cli_3.6.2
## [43] rmarkdown_2.23               crayon_1.5.2
## [45] generics_0.1.3               rstudioapi_0.15.0
## [47] rjson_0.2.21                 ggforce_0.4.2
## [49] stringr_1.5.1                zlibbioc_1.48.2
## [51] parallel_4.3.1               BiocManager_1.30.20
## [53] XVector_0.42.0               restfulr_0.0.15
## [55] matrixStats_1.3.0            vctrs_0.6.5
## [57] V8_4.4.2                     Matrix_1.6-3
## [59] jsonlite_1.8.8               bookdown_0.34
## [61] IRanges_2.36.0               S4Vectors_0.40.2
## [63] gggenes_0.5.1                randomcoloR_1.1.0.1
## [65] magick_2.8.1                 ggnewscale_0.4.10
## [67] wesanderson_0.3.7            glue_1.7.0
## [69] codetools_0.2-19             stringi_1.8.4
## [71] gtable_0.3.5                 GenomeInfoDb_1.38.8
## [73] BiocIO_1.12.0                GenomicRanges_1.54.1
## [75] munsell_0.5.1                tibble_3.2.1
## [77] pillar_1.9.0                 htmltools_0.5.5
## [79] ggfittext_0.10.2             GenomeInfoDbData_1.2.11
## [81] BSgenome_1.70.2              R6_2.5.1
## [83] evaluate_0.21                lattice_0.21-8
## [85] Biobase_2.62.0               Rsamtools_2.18.0
## [87] Rcpp_1.0.12                  SparseArray_1.2.4
## [89] xfun_0.44                    MatrixGenerics_1.14.0
## [91] pkgconfig_2.0.3
```
